# Lipid membranes modulate the activity of RNA through sequence-dependent interactions

**DOI:** 10.1101/2021.03.25.437010

**Authors:** Tomasz Czerniak, James P Saenz

## Abstract

RNA is a ubiquitous biomolecule that can serve as both catalyst and information carrier. Understanding how RNA bioactivity is controlled is crucial for elucidating its physiological roles and potential applications in synthetic biology. Here we show that lipid membranes can act as RNA organization platforms, introducing a novel mechanism for ribo-regulation. The activity of R3C ribozyme can be modified by the presence of lipid membranes, with direct RNA-lipid interactions dependent on RNA nucleotide content, base pairing and length. In particular, the presence of guanine in short RNAs is crucial for RNA-lipid interactions, and G-quadruplex formation further promotes lipid binding. Lastly, by artificially modifying the R3C substrate sequence to enhance membrane binding we generated a lipid-sensitive ribozyme reaction with riboswitch-like behavior. These findings introduce RNA-lipid interactions as a tool for developing synthetic riboswitches and novel RNA-based lipid biosensors, and bear significant implications for RNA World scenarios for the origin of life.

## Introduction

RNA performs diverse functions ranging from information storage to regulation of other biomolecules and direct catalysis of biochemical reactions. The functional versatility of RNA has implications for understanding plausible scenarios for the origin of life^1–3^, and for developing tools in synthetic biology^4–6^. Research aimed at understanding how an RNA World could have emerged has motivated development of ribozymes with functions including RNA ligation^7–10^, replication^11,12^ and other activities^13–16^. The experimental development of functional RNAs raises the possibility of recapitulating an RNA World and engineering biochemical systems based on RNA. For both synthetic biology and understanding the origin of self-replicating organisms, RNA has intrinsic appeal: it can serve functions of both DNA (information storage) and proteins (enzymes), obviating the need for translation machineries and protein chaperones. Furthermore, RNAs are more likely than proteins to undergo some degree of reversible denaturation to a functional conformation, lending robustness against a broad range of physical and chemical conditions. In order to design a biochemical system based on RNA, however, it is essential to be able to coordinate RNA activity in space and time.

Key to harnessing the functional versatility of RNA is understanding how to spatially and temporally modulate its properties and to selectively modulate the activity of different RNAs within one system. The physicochemical environment surrounding an RNA molecule is a central determinant of its structure, stability and activity. Spontaneous RNA hydrolysis and ligation, as well as catalytic RNA activity are sensitive to pH^17^, ionic strength^18,19^ and RNA concentration changes^11^, among other parameters. Similarly, RNA activity can be modulated by interactions with molecules such as ions, proteins, and other nucleic acids^20^. Thus, one approach to regulating RNA activity could be via tunable interactions with binding partners that affect RNA structure, concentration or chemical microenvironment.

One mechanism for modulating RNA activity could be through direct RNA-lipid interactions^19,21,22^. Because of their amphiphilic nature, lipids spontaneously self-assemble into membranous structures that can encapsulate RNA into protected and selective microcompartments^23–25^. Alternatively, direct RNA-lipid interactions could localize RNAs to membrane surfaces, increasing its local concentration and reducing dimensionality for intermolecular interactions^21^. Lastly, localization to a lipid surface brings RNA into a physicochemically unique microenvironment with sharp gradients of hydrophobicity, electrical permittivity, and water activity. Through these effects, RNA-lipid interactions could provide a powerful mechanism for modulating RNA activity.

The first functional RNA-lipid interaction was described more than 40 years ago^26^, with subsequent research revealing various factors that facilitate nucleic acid-lipid binding^27–37^. More recently, specific RNA sequences have been generated through SELEX with affinity for fluid membranes comprised of phospholipids and cholesterol^38–40^. Interestingly, mixtures of RNAs have also been shown to bind to membranes that are in a solid crystalline (gel) phase^41,42^. These studies revealed that, while most randomized mixtures of RNA sequences can bind to gel membranes, there is a relatively small chemical space of oligomers that have affinity for fluid membranes. Thus, conceptually, gel membranes could provide a platform for modulating the activity of a diverse range of RNAs. However, the effects of gel membranes on RNA activity and the sequence selectivity of such interactions are relatively unexplored.

This study reports the effect of lipid membranes on RNA catalytic activity. We show that RNA-lipid binding depends on the primary sequence, secondary structure, and length of RNA. Using the transacting R3C ligase ribozyme, we observed that R3C-lipid binding changes ribozyme activity in a concentration-dependent manner. Lipid binding assays show that the interaction of short RNA sequences with gel membranes depends on guanine content and the presence of double-stranded structures. Lastly, modification of R3C’s substrate sequence increased the tunability of R3C-based reactions through a lipid-dependent mechanism. Our findings demonstrate that membranes can serve as platforms for ribo-regulation, which could contribute to the development of RNA-based lipid biosensors and lipid-sensitive riboswitches. This approach introduces new tools for molecular and synthetic biology and raises the prospect of previously unrecognized roles for RNA-lipid interactions in the origin and evolution of life.

## Results

The discovery that RNA can catalyse reactions in addition to encoding information^18,43^, opened new directions for engineering life and the possibility of protocells emerging from an RNA world^2^. But, a key missing ingredient for RNA-based systems (e.g. an RNA World or synthetic systems based on RNA biochemistry) is a mechanism to organize RNAs and regulate their activity. We hypothesized that RNA-membrane interactions could influence ribozyme activity by changing local RNA concentrations at the membrane surface or influencing RNA conformations.

We took advantage of the observation that RNA can interact with solid crystalline (gel) phase membranes composed of phosphatidylcholine lipids^41,42^ to test the hypothesis that lipid membranes can serve as platforms for ribo-regulation. We first determined the membrane-buffer partition coefficients for a random mixture of RNA oligomers using phosphatidylcholine lipids employed in previous work^42^ that are in a gel phase (DPPC at 24 °C), ripple phase (DMPC at 24 °C), a liquid disordered (L_d_) phase (DOPC) and a liquid disordered-liquid ordered (L_d_-L_o_) phase separated system (DOPC:DPPC:cholesterol, 2:2:1 ratio) (**Fig. 1**). RNA lipid-buffer partition coefficients indicate how well a particular molecule binds to the membrane by comparing the relative amount of RNA in buffer and on the lipid membrane^44^. As expected, RNA showed the strongest binding to gel phase membranes with a greater than 10-fold higher partition coefficient than for fluid membranes. Interestingly RNA showed slightly higher binding to L_d_-L_o_ phase separated membranes than to L_d_ phase membranes, consistent with previous observations indicating that RNA can have a higher affinity for membranes in the more rigid L_o_ phase^42^. Surprisingly, RNA bound comparatively well to gel membranes composed of the saturated fatty acid palmitate, with a partition coefficient falling in between fluid and gel phosphatidylcholine membranes. Fatty acids are among the simple amphiphiles that could have accumulated on early Earth^45^. Thus, RNA-lipid interactions can now be extended to a prebiotically plausible lipid. It is worth noting that most of the RNA oligomers measured in this study had partition coefficients above 10^5^, which is in the range of partition coefficients determined for hydrophobic peptides^44^. Binding to DPPC membranes remained above 10^5^ even at physiological concentrations of Ca and Mg ions (**Suppl. 1a**). These observations allowed us to identify DPPC gel membranes as the most optimal lipid for this study, based on the superior RNA-lipid binding coefficient for DPPC.

**Fig. 1.**
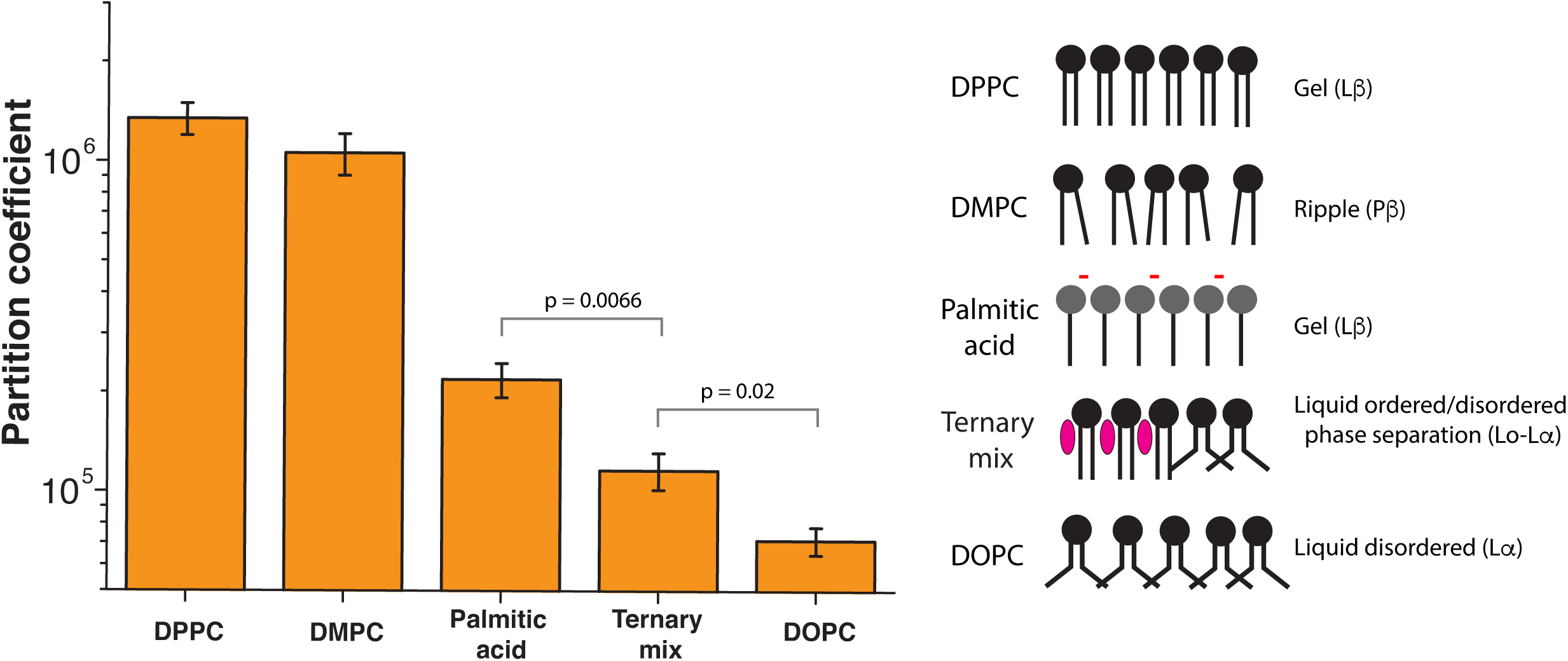
RNA-lipid binding depends on membrane fluidity. Lipid-buffer partition coefficients were determined for the 40 nt random RNA sequences with phosphatidylcholine and fatty acid vesicles. The highest binding was observed for phospholipid-based gel membranes (DPPC, partition coefficient >1×106) and ripple-phase membrane (DMPC at 24 °C, partition coefficient of 1×106) whereas the lowest binding was observed for liquid disordered membranes (DOPC, partition coefficient <1×105). The partition coefficient for the palmitic acid gel membranes is significantly higher than for the fluid ternary mixture and DOPC membranes.

To visibly demonstrate the preferential binding of RNA to gel versus fluid membranes, we prepared giant unilamellar vesicles (GUVs) that are phase separated into gel and liquid domains and observed the distribution of a random mixture of RNA oligomers stained with SybrGold by fluorescence microscopy (**Fig. 2a**). A fluorescent lipid probe (DiD) was enriched in membrane liquid domains and was excluded from gel phase domains, which allowed us to directly observe specific RNA-gel membrane colocalization in a phase separated system. RNA enrichment on GUV gel domains was reversed by heating the system above the gel phase melting temperature, resulting in an entirely liquid phase vesicle (**Fig. 2b**). Binding of RNA oligonucleotides to gel-phase small unilamellar vesicles (SUVs) can also lead to aggregation into large visible RNA-lipid assemblies^37,41^ (**Fig. 3a**), which is probably due to charge-based interactions (**Suppl. 1c**). Depletion of divalent cations (both Mg and Ca, **Suppl. 1b**) or increasing temperature above the lipid melting temperature (i.e. producing fluid instead of gel membranes) reduced RNA-membrane binding and aggregation (**Fig. 3a**)^27,31,41,42^. Taken together, reversible RNA-lipid binding and lipid-dependent aggregation could provide a tunable mechanism to concentrate and regulate RNAs in simple bottom-up synthetic systems, or in a prebiotic environment.

**Fig. 2.**
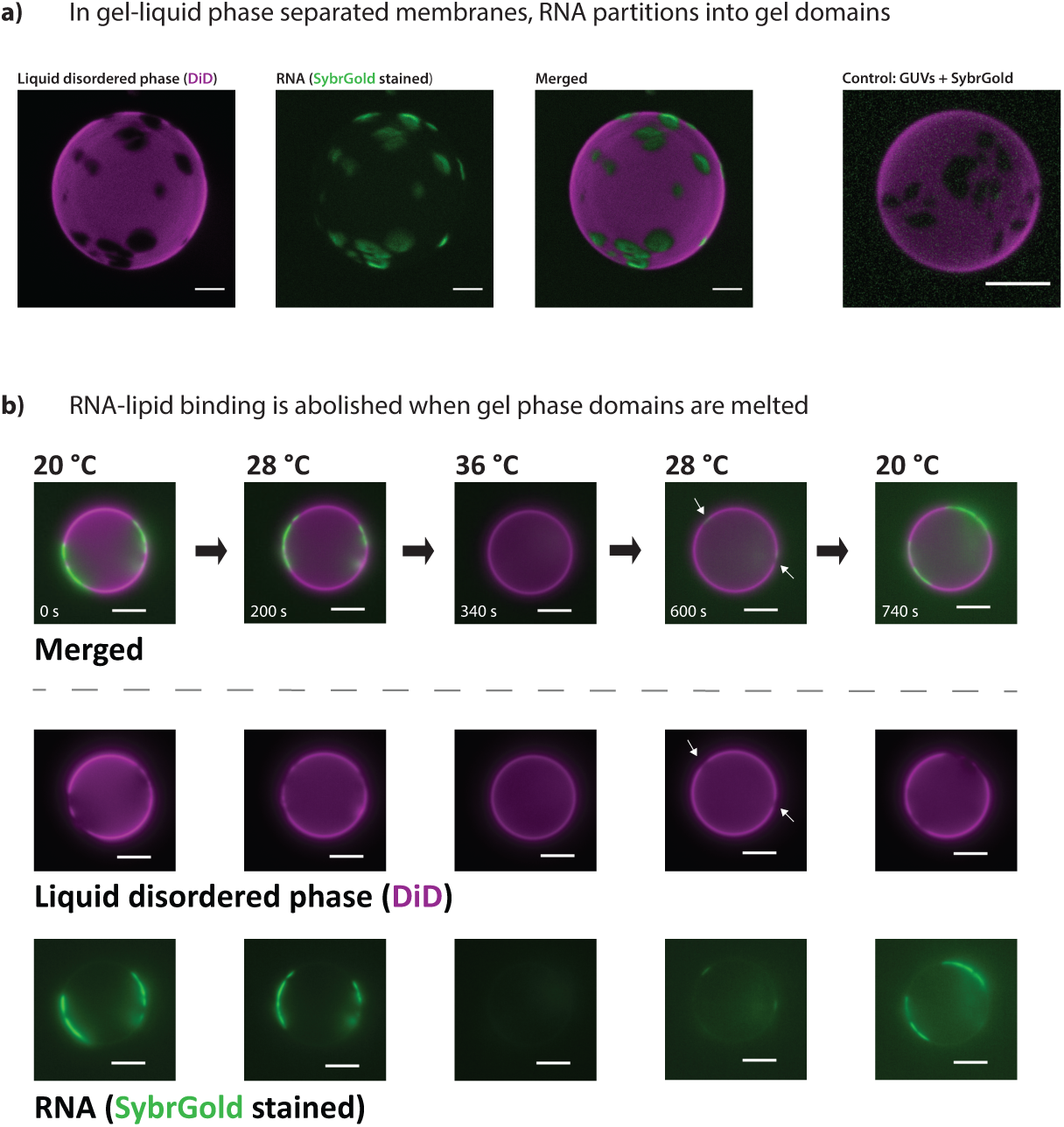
RNA selectively binds to gel phase membrane domains. **(a)** Gel-liquid phase separated giant unilamellar vesicles (GUVs) were prepared from a mixture of DOPC and DPPC with a molar ratio of 1:1 and labeled with 0.5 mol% of the fluorescent lipophilic probe DiD. DiD is excluded from gel phase domains, which are observed as non-stained regions on the surface of the vesicle. A mixture of random RNA oligomers (40xN) stained with SybrGold is enriched within the gel phase domains. A control without RNA shows that SybrGold itself does not stain the GUVs. **(b)** When gel phase domains are melted by increasing temperature (3.4 °C/min – see **supplementary movie 1**) RNA no longer enriches at the GUV surface. Decreasing the temperature again restores gel-liquid phase separation and, consequently, RNA-membrane binding (∼28 °C, white arrows – see **supplementary movie 2**). All of the scale bars are 5 μm.

**Fig. 3.**
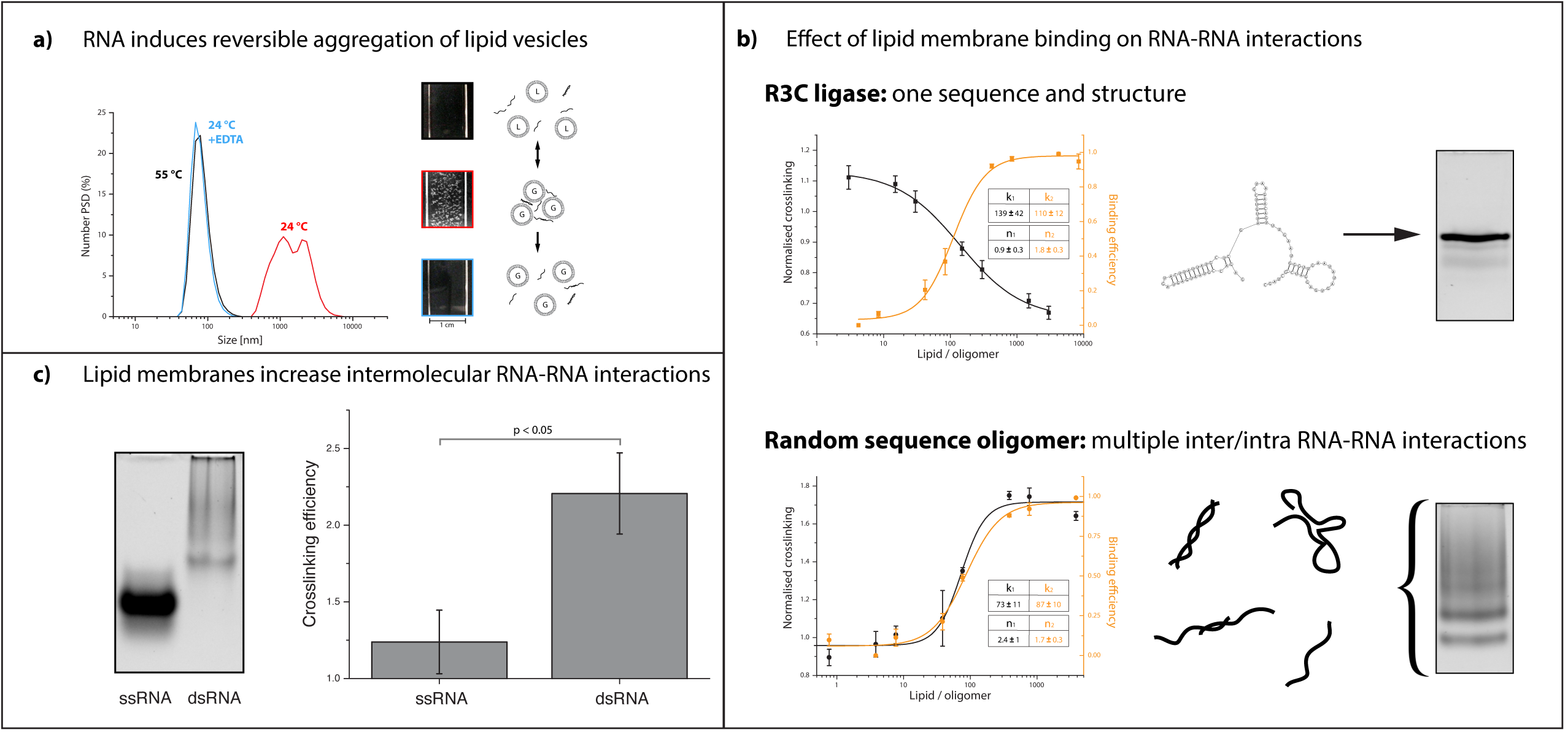
RNA-lipid interactions induce reversible aggregation of vesicles and enhance RNA-RNA interactions. **(a)** RNA-dependent lipid vesicle aggregation can be observed through changes in vesicle size distribution measured by dynamic light scattering (left), or visually as depicted in the cuvette images (right). The introduction of a randomized pool of RNA oligos to gel lipid membranes (gel phase = G) induces vesicle aggregation, which can be reversed either through chelation of divalent cations with EDTA or by increasing the temperature to achieve membranes in a liquid phase (liquid phase = L). **(b)** R3C ligase (84 nt), which is represented by one structure (see right, native PAGE) shows a small increase in UV-crosslinking at low lipid concentration, whereas crosslinking efficiency decreases at higher lipid concentration. Binding and UV crosslinking are inversely correlated. Randomized RNA oligos (75-85 nt, mix of different inter- and intramolecular structures) show increasing crosslinking efficiency in the presence of lipids; crosslinking and lipid binding are closely correlated. Both R3C and randomer mix have a similar size range (**Suppl.2**). All of the fits are Hill’s function fits; errors are standard error of the fit based on at least 3 replicates. **(c)** The ratio of UV-crosslinking efficiency with and without lipid vesicles was calculated separately for single stranded RNA and double stranded RNA - larger values indicate a higher degree of lipid-dependent crosslinking. Double stranded RNA crosslinking efficiency is enhanced by the presence of lipid membranes, whereas a smaller effect is observed for single stranded RNAs.

### RNA-lipid binding changes the probability of RNA-RNA interactions

To determine whether local RNA concentration is influenced by interaction with lipid membranes, we relied on UV-mediated crosslinking. All small vesicle membranes in this study were composed of DPPC, which is in a solid crystalline (gel) phase at the experimental temperature of 24°C. Nucleic acid bases absorb UV light, producing chemical changes that yield base-base covalent bonds in a distance-dependent manner, yielding insights into the structure and interactions of nucleic acids^46–48^. We first observed the effect of gel membranes on the crosslinking of a single defined RNA sequence, the R3C ligase, which predominately forms one structure (**Fig. 3b, upper native gel**). In subsequent experiments, we focus on the R3C ligase, since it is a relatively short single turnover ribozyme with a simple structure and a prebiotically interesting ligation activity (**Suppl. table 1**)^10^. When subjected to UV, the R3C ligase shows slightly increased crosslinking in the presence of low lipid concentrations. A further increase in lipid concentration led to decreased crosslinking efficiency, possibly through dilution of the RNA species on the surface of the lipid membranes. This effect was inversely correlated with RNA-lipid binding efficiency (**Fig. 3b**). In contrast, a mixture of RNA oligomers with randomized sequences which can form a more diverse range of inter- and intra-molecular structures (**Fig. 3b, bottom native gel**) showed continuously increasing and overall higher UV-crosslinking efficiency in the presence of lipids (**Fig. 3b, bottom graph**). These results show that RNA-gel membrane binding can influence the local concentration of RNAs in very different ways depending on the type of RNA.

The contrasting effect of crosslinking for R3C and randomized oligomers suggested that gel membrane binding can enhance RNA-RNA interactions for RNAs with a higher propensity for intermolecular interactions. Indeed, we observed that a mixture of two oligomers with complementary sequences showed enhanced crosslinking relative to a single oligo with lower propensity for inter- and intramolecular interactions (**Fig. 3c**). At higher lipid:RNA ratios two things happen: a larger fraction of the total RNA becomes bound to the membrane, and available membrane surface area increases thereby diluting the lateral density of RNAs on the membrane. Increased RNA crosslinking for randomized oligomers at higher lipid:RNA ratios is therefore most probably influenced by an enhancing effect of membrane binding on RNA-RNA interactions, which becomes more prominent as a larger fraction of the total RNA is bound to the membrane surface. Thus, RNA-membrane interactions can influence RNA-RNA interactions in a manner that is dependent on lipid concentration and RNA sequence diversity. This further suggests that membrane binding could have an effect on trans-acting ribozyme activity derived from base pairing (**Suppl. table 1**).

### RNA-lipid interactions influence RNA catalytic activity

To investigate the functional consequence of RNA-lipid binding, we tested the effect of lipids on the activity of the trans-acting R3C ligase ribozyme. R3C ligase is a ribozyme that catalyzes ligation of substrate strands to the ribozyme (**Fig. 4a**)^10^. Since R3C is also part of an RNA self-replication system, the effects of lipids on R3C are also interesting with regard to the emergence, evolution and artificial synthesis of autonomous self-replicating systems^11,12^. The R3C reaction rate in the absence of membranes was 11 pM/min, with a reaction constant rate of k = 6.4×10^−4^ min^-1^. Addition of lipid vesicles led to an increased reaction rate (14 pM/min, +29%), which could plausibly be due to increased ligase-substrate interaction on the membrane either through increased concentrations at the membrane surface or through enhanced exposure of the substrate-binding domain of R3C. At the highest lipid concentration, the reaction rate dropped to 8 pM/min (−27%) (**Fig. 4b**). A decrease in reaction rate was correlated with further R3C ligase binding to the lipid membranes (>100 lipid/R3C, **Fig. 3b**). We speculate that this may be caused by dilution of RNA on the membrane as the available membrane surface area increases at higher lipid:RNA ratios. Alternatively, if preferential membrane binding of either R3C or its substrate occurs, then aggregation of lipid vesicles could selectively sequester RNA within vesicle aggregates reducing interactions of the ribozyme with its substrate. Importantly, we did not observe enhanced activity relative to the vesicle-free reaction in the presence of fluid membranes composed of DOPC, or gel membranes in the absence of Ca ions (**Suppl. 3**), both conditions in which RNA-lipid binding is impaired. Thus, despite the relatively small effect on activity, these results provide the proof of principle that RNA-gel membrane binding can enhance the reaction rate of a trans-acting ribozyme in a lipid concentration-dependent manner.

**Fig. 4.**
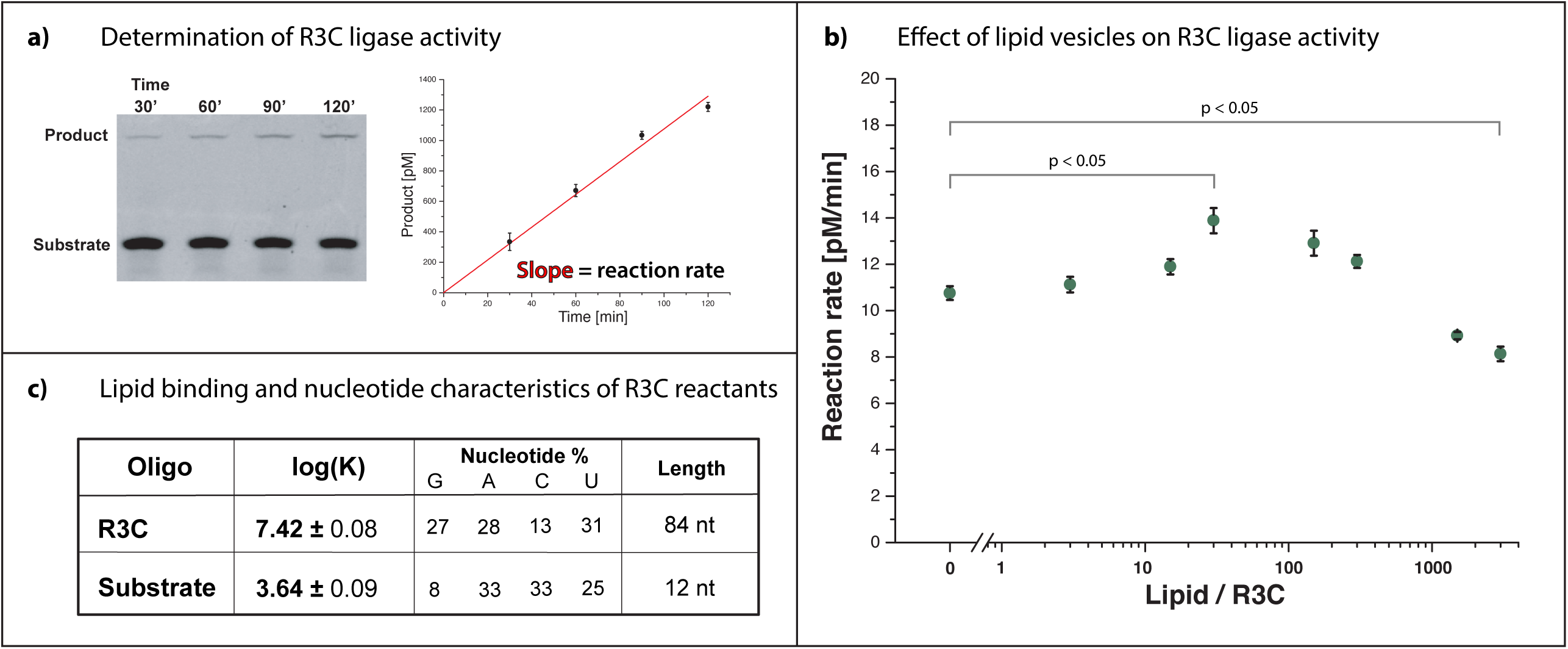
Lipid membranes influence R3C ligase activity through lipid-ligase interactions. **(a)** R3C ligase reaction rates were quantified by measuring the amount of product within a two hour incubation time. **(b)** R3C reaction rate is dependent on the presence of gel membranes - increased activity is observed for lower lipid concentration followed by a drop in activity at higher lipid concentration. Error bars represent standard errors from concatenate linear fits from 3 replicates. **(c)** The 12 nt R3C substrate shows several orders of magnitude lower membrane-buffer partition coefficient log(K) compared with the R3C ligase, which could be derived from different compositional or structural characteristics of both RNA species. Error ranges represent standard error calculated from at least 3 binding replicates.

### R3C ligase and its substrate bind differently to the membrane

To further understand if preferential membrane binding of either R3C or its substrate contributes to changes in ligation rates, we analyzed lipid-binding affinities of the ligase and its substrate. The partition coefficient of the short 12 nt substrate was more than 3 orders of magnitude lower than that of the 84 nt R3C ligase (**Fig. 4c**). Thus, in this system, substrate-membrane binding is essentially negligible compared with R3C-membrane binding, and this could partly explain why we observed such a small (∼30%) enhancement in R3C ligation activity. The negligible substrate binding suggests that enhanced activity could be due to an effect of R3C-lipid interaction on the ligase-substrate complex, consistent with our observation that the probability of RNA-RNA interactions can be enhanced through RNA-lipid binding (**Fig. 3c**). Alternatively, enhanced activity could be derived from interactions of the substrate with the membrane after it has bound to R3C. Such a large difference in binding was surprising since we had begun this study with the assumption based on previous observations that RNA-gel membrane binding is far less selective than for RNA-fluid membrane binding^42^. The most prominent differences between R3C and its substrate are length and nucleotide composition (**Fig. 4c**), and this prompted us to explore which features of RNA could influence binding to gel membranes.

### RNA sequence influences binding to gel membranes

Understanding the features that influence RNA-lipid interactions could allow us to engineer RNAs with higher membrane affinity, and thereby enhance activity by concentrating RNAs at the membrane surface. The interaction of RNA with fluid membranes has been shown to be very specific to sequence and structure^38–40^. In contrast, interactions of RNA with gel-phase membranes have been previously shown to exhibit low sequence-dependence, based on binding of oligomers with various RNA sequences^41,42^. However, the large difference in binding for R3C and its substrate suggests that RNA-gel membrane interactions might in fact be dependent on RNA composition or structure.

To investigate which features of RNA influence its interaction with lipid membranes we first evaluated how base content influences membrane binding of short RNA species. 19 nt oligomers with only one type of nucleotide were tested together with a random sequence as a control. Remarkably, whereas the 19xG oligomer bound very efficiently to the gel-phase membrane (partition coefficient >1×10^6^), oligomers of A, C or U showed practically no binding. A mixture of 19 nt RNA with randomized sequences showed moderate binding, as did one with AG-repeats (**Fig. 5a**), most probably due to the introduction of guanine. While the formation of structures through non-canonical A-G pairing^49–51^ could potentially influence binding, migration of the AG-repeat oligomer as a single band on non-denaturing gel argues against that possibility (**Fig. 7b**). Enhanced binding of an oligomer with only AG-repeats, therefore, suggests that membrane binding can be influenced by direct guanosine-membrane interactions^52^. In general, guanine could conceivably enhance binding either through direct interactions with the membrane, or alternatively by promoting G≡C base pairing or presence of G-quadruplexes, thereby influencing structure.

**Fig. 5.**
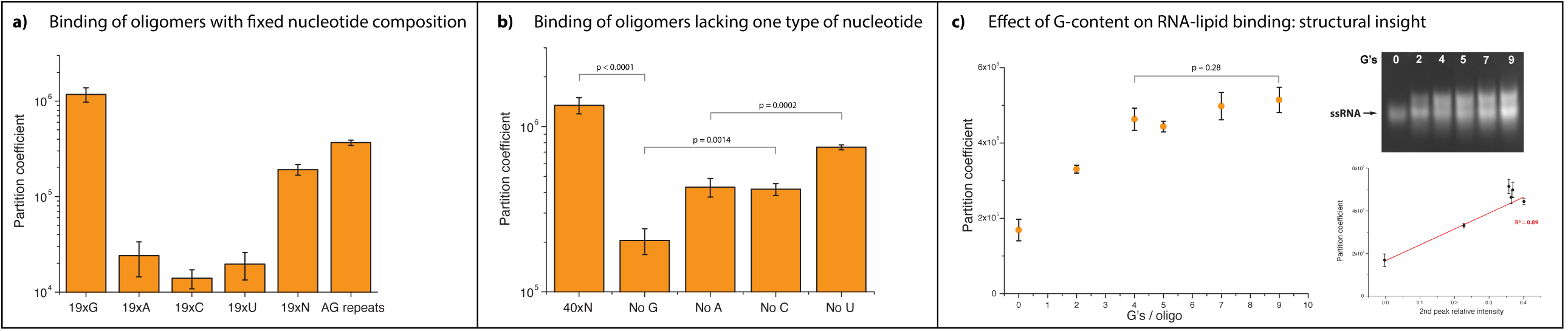
RNA-lipid binding is RNA sequence-dependent. **(a)** Short 19 nucleotide oligomers show nucleotide-dependent binding to gel membranes, with highest binding observed for the pure G oligomer followed by oligos with randomized sequence or AG repeats. Oligomers containing only A, C or U bases show negligible binding to lipid membranes, compared with the pure G oligomer (p value < 0.05). **(b)** Oligonucleotide length also influences binding efficiency as seen in the comparison of **(a)** 19xN and **(b)** 40xN. Depletion of one type of nucleotide within oligomers lowers binding efficiency. The absence of G decreased partition coefficient by 6.6x compared to non-depleted RNA, whereas absence of A, U or C showed a less pronounced effect. Significant differences are present between G- and C-deficient RNAs, as well as between A- and U-deficient RNAs. **(c)** Oligonucleotides with increasing G content showed higher lipid – buffer partition coefficients with values plateauing for oligomers containing more than four G’s. Increased binding is correlated with the appearance of second band on native gel, which suggests that structural changes within RNA oligomers might contribute to lipid membrane binding.

We next examined the effect of deleting a specific base from mixtures of 40 nt oligomers with randomized sequences. The depletion of G led to the largest decrease in binding, followed by an intermediate decrease in binding from depletion of A and C, and the smallest decrease in binding from depletion of U. The decrease in binding affinity in all of the base-depleted oligomers (**Fig. 5b**) indicates that base-pairing may influence membrane binding. However, the larger effect of G depletion than C depletion indicates that base pairing is not the only factor responsible for membrane binding, consistent with our previous observation that G-content alone can influence binding (**Fig. 5a**). The depletion of A also generates a larger effect than U depletion, suggesting that other factors such as non-canonical base pairing could play a role. Finally, by comparing 19 nt and 40 nt randomized oligomers, we observed that increasing RNA length also led to enhanced binding, possibly due to increased length and G content (**Fig. 5a, b**).

To further determine how guanine content affects membrane binding, we measured lipid-buffer partition coefficients for short oligomers with varying G-content. Remarkably, addition of two G’s per oligomer led to a two-fold increase of the partition coefficient relative to guanine-depleted RNA. Oligomers with four and more guanine residues showed a plateau of partition coefficient values (**Fig. 5c**). The saturable effect of increasing G-content indicates that binding is not solely due to cumulative guanine-lipid interactions, but also to other attributes related to G-content, such as base paring-derived structures. Analysis by non-denaturing gel shows that a second band appears with the introduction of G, and that its relative intensity correlates with binding (**Fig. 5c**), suggesting that the formation of G-dependent intra- or intermolecular structures influences RNA-lipid interactions. Indeed, tuning the distribution and spacing of G in oligomers with fixed 2xG content resulted in varying binding that was also correlated with the appearance of a second band on non-denaturing gel (**Suppl. 4**). These results indicate that G-dependent structures influence RNA-lipid binding and that even low guanine content can significantly increase membrane binding efficiency (**Fig. 5c**).

### RNA-membrane binding is dependent on RNA base pairing

The effect of base depletion on oligomer binding suggested that base-pairing may be a key factor in membrane binding. To investigate the effect of base-pairing on membrane interactions, we estimated binding efficiency for 36 nt ssRNA and dsRNA (composed of complementary CAGU and ACUG repeats). We observed that not only was the membrane binding efficiency higher for dsRNA compared with ssRNA species, but also that only dsRNA promoted vesicle aggregation (**Fig. 6a**). Binding of dsRNA with a non-repetitive sequence was also relatively high (partition coefficient of 4×10^6^, **Suppl. 5**), controlling for the possibility that higher binding of dsRNA was due to longer intermolecular linkages formed by staggering of repeated complimentary ACUG/CAGU strands. We further observed that dsRNA interacts with fewer lipids than ssRNA, implying different models of binding (**Fig. 6b**). Thus, membrane binding efficiency can be dependent on the propensity of an RNA molecule to form intra- or inter-molecular structures through base pairing. Since base pairing is a fundamental element of RNA secondary structure, our results indicate that structure in general can influence the selectivity of RNA-lipid interactions.

**Fig. 6.**
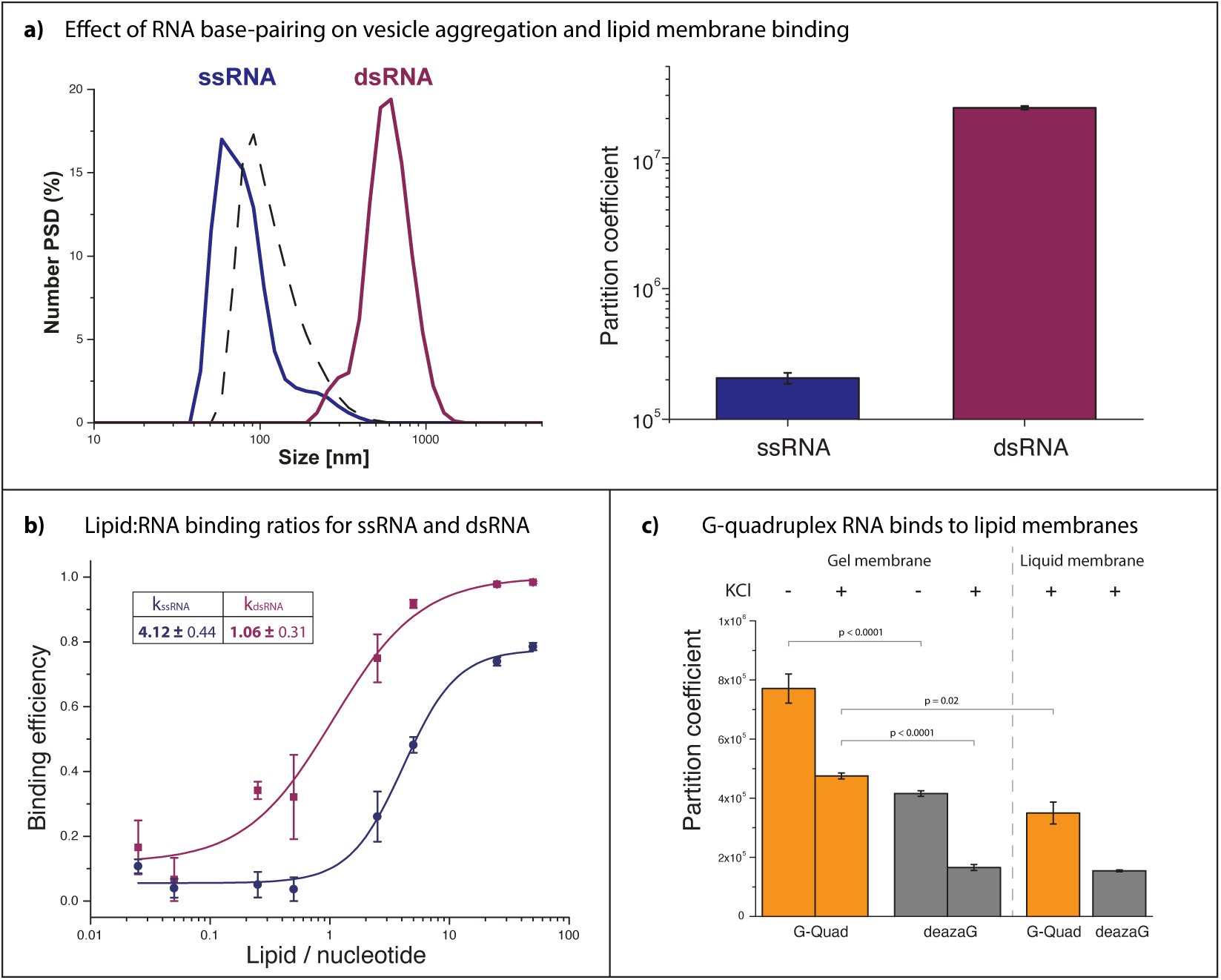
RNA-gel membrane binding depends on RNA structure. **(a)** The effect of RNA base-pairing structure on lipid vesicle aggregation (left) and membrane binding (right) was evaluated by comparing RNAs with limited ability to create inter- and intramolecular structures (ssRNA: 36 nucleotide CAGU repeats) with 100% complementary double stranded RNAs (dsRNA: mixed ACUG and CAGU oligos). Single stranded RNAs do not cause lipid vesicle aggregation despite moderate membrane binding, whereas double stranded RNA species show both high binding and vesicle aggregation. Dashed black line: DPPC vesicles without RNA. **(b)** RNA:lipid binding ratios (shown in table) estimated from binding curves of ssRNA and dsRNA (plotted) are affected by base pairing. **(c)** 19 nt G-quadruplex forming RNA binds to both gel and liquid membranes in the presence and absence of KCl ions. The G-quadruplex oligomer has significantly higher partition coefficients than an oligo with the same sequence containing deaza-guanine, which can’t form G-quadruplex structures.

### RNA G-quadruplex structures bind well to both gel and fluid membranes

The correlation between G content and enhanced membrane binding pointed towards a potential role for G-dependent RNA structures. G-rich RNAs can form structures known as G-quadruplexes, in which G-G interactions lead to the formation of stacked G-tetrads^53^. We therefore examined whether G-quadruplex formation enhances RNA-lipid interactions. To do this, we synthesized a G-quadruplex forming RNA with 7-deaza-guanine instead of guanine to inhibit G-quadruplex formation^54^. The guanine containing oligomer showed almost 2-fold higher binding coefficient than the 7-deaza-guanine containing oligomer. Although G-quadruplexes can exist in the presence of divalent ions^55^, they are most effectively stabilized by potassium ions, which was absent from the buffer. However, salts composed of monovalent ions like KCl can inhibit RNA-lipid binding^41,42^. Consequently, addition of potassium ions to the buffer led to an overall decrease in binding, but the relative difference in binding between guanine- and 7-deaza-guanine-containing oligomers was increased to almost 3-fold, demonstrating that G-quadruplex formation enhances gel membrane binding (**Fig. 6c**).

Since G-quadruplexes are physiologically relevant structures with diverse roles in cellular regulation^56^, we asked whether they might also show enhanced binding to fluid physiologically relevant membranes. Much to our surprise, the partition coefficient for our G-quadruplex oligomer for fluid membranes composed of DOPC was within the same range as partition coefficient measured for gel membranes (**Fig. 6c**).

### Modifying R3C substrate sequence enhances membrane binding and modifies reaction rates

In previous experiments, we observed that the binding affinity of the short substrate was >3 orders of magnitude lower than for the R3C ligase and that reaction rates correlated with R3C-membrane binding (**Fig. 4b, c**). We therefore reasoned that by modifying the R3C ligase substrate to enhance membrane binding, we might be able to further modulate ligase activity through increased RNA density at the membrane surface. Having determined that guanine content is a key factor influencing RNA-lipid binding (**Fig. 5**), we modified the R3C substrate sequence to increase lipid membrane binding efficiency by addition of a 5’-overhang with varying amounts of AG repeats (**Fig. 7a**).

**Fig. 7.**
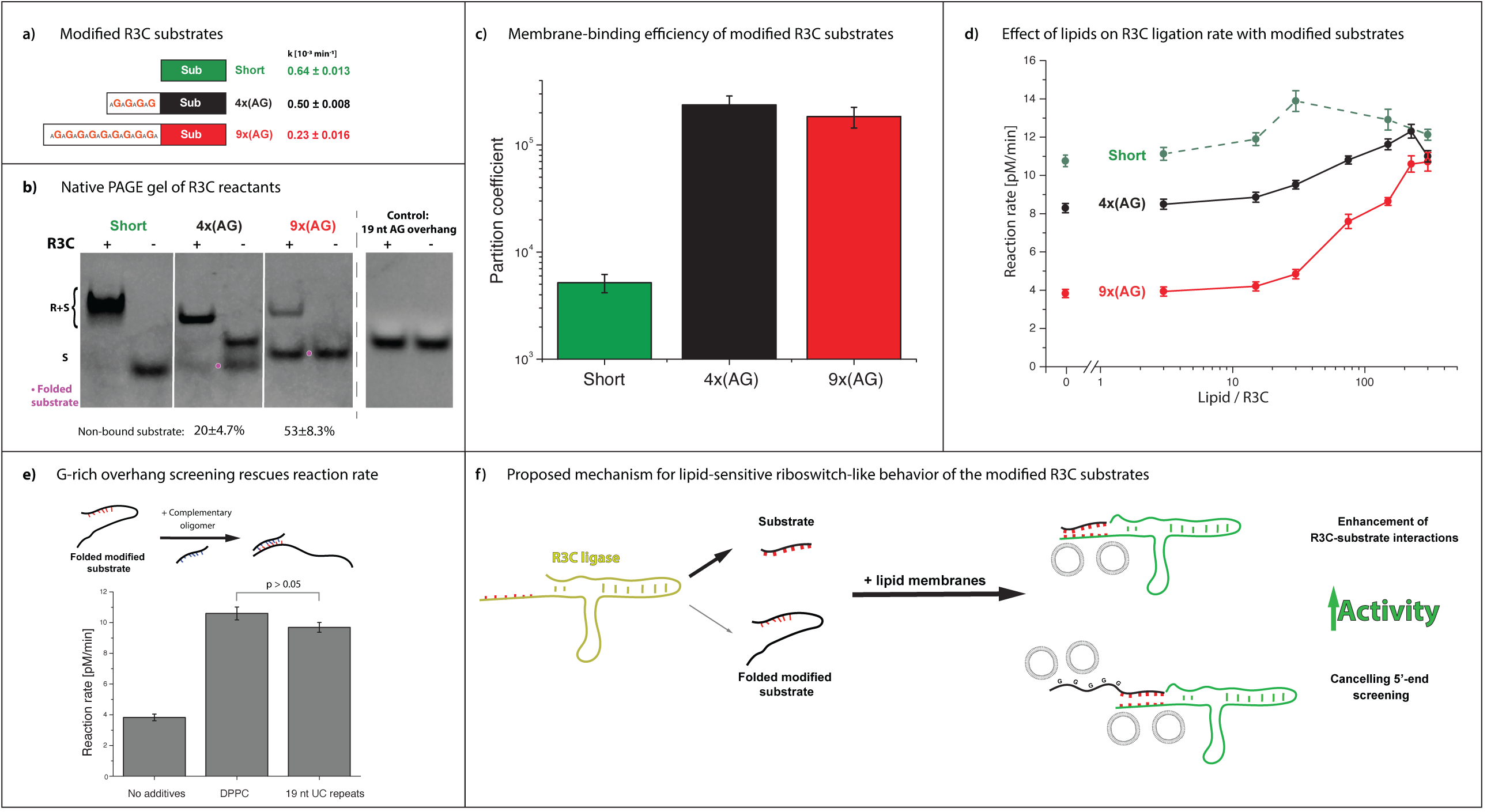
Modification of the R3C substrate sequences enables lipid-dependent tunability of ligation rate. **(a)** Modified R3C reaction substrates were synthesized in order to increase binding to the membrane. The catalytic 3’ end of the molecule was enriched with 5’ tails with varying G content. Reactions for modified substrates showed lower reaction rate constants compared with the unmodified substrate. **(b)** The mechanism for inhibition of activity was evaluated by analysis of R3C reactants using native PAGE. The unmodified short substrate is entirely bound to the ligase, whereas AG-modified substrates do not bind entirely and show mobility abnormalities, which suggest substrate folding (magenta points). An AG-rich oligomer does not fold into multiple structures itself and does not comigrate with the ligase, indicating that folding of the substrates impairs substrate-ligase interactions. **(c)** Modification of the substrate sequence through addition of a 5’ overhang increases binding to the membrane. **(d)** Addition of lipid membranes to reactions with the G-rich substrates increases reaction rates to the level of the unmodified substrate. **(e)** Addition of an oligomer with complementary sequence to the 5’end of 9x(AG) substrate overhang rescues R3C activity, comparable to the effect of gel membranes. **(f)** We propose that the decrease of R3C reaction rates in the presence of G-rich substrates is based on the inhibition of the catalytic part of the substrate by the 5’ G-rich overhang. The presence of lipid membranes not only increases R3C-substrate interactions, but also screens the 5’ G-rich substrate part enabling the R3C reaction to proceed.

We first assessed the effect of modifying the R3C ligase substrate on ligation rate in the absence of membranes (**Fig. 7a**). We observed that increasing the guanine content decreased ligase activity, suggesting that 5’ modification of the substrate was inhibiting the reaction either through steric hindrance or promotion of inhibitory inter- or intra-molecular interactions. To determine whether AG-rich substrates form intra-molecular structures or exhibit reduced binding with the R3C ligase, we performed electrophoresis in native conditions with and without ligase (**Fig. 7b**). Firstly, the non-modified short substrate migrated as one band and was fully bound to the R3C ligase. Secondly, a 4x(AG) 20 nt substrate variant migrated as two bands, which suggests that part of the substrate molecules have some folding that might bias binding to the R3C ligase. In the presence of ligase, 20 ± 4.7% of the substrate was non-bound, which correlated with a 22% reduction in reaction rate. For the longest substrate variant (31 nt), electrophoretic mobility suggested higher folding, since only one band was present and migrated half-way between the non-folded 12 nt and 20 nt substrates. 53 ± 8.3% of the 9x(AG) substrate was not bound to the ligase, which correlates with a reduced R3C reaction rate. Lastly, a free AG-rich 19 nt oligomer did not co-migrate with the R3C ligase, indicating that the inhibition effect is not based on interactions between the G-containing substrate overhangs and the ligase (**Fig. 7b**). We further observed that 5’ modification of the substrate with a G-depleted randomized 19 nt overhang had an insignificant effect on R3C activity, and modification with a 2xG randomized 19 nt overhang produced a relatively small effect (**Suppl. 6**). Therefore, increasing guanine content reduces R3C ligase-substrate binding, likely accounting for decreased activity.

We next investigated whether varying guanine content through 5’ modification of the substrate influences membrane binding and, consequently, ligation activity in the presence of membranes. As expected, the longer guanine-containing R3C substrates showed significantly higher membrane binding compared with the shorter variant (**Fig. 7c**). To determine whether binding of the substrates to the membrane enhances ligase activity, we determined R3C reaction rates in the presence of the gel lipid membranes (**Fig. 7d**). For modified substrates, the reaction rates increased significantly to the level of the unmodified substrate. Interestingly, none of the modified substrates exhibited higher activity than the unmodified substrate, indicating that the rescue of activity of the other guanine variants is not due to increased substrate-ligase interactions through enhanced binding to the membrane, but rather due to interaction of the membrane with 5’ overhangs. We confirmed that the effect on activity is due to membrane-substrate overhang interactions by showing a rescue of ligation activity upon the addition of an oligomer with a complimentary sequence to the 5’ substrate modifications (**Fig. 7e**). Thus, substrate-membrane interaction increases activity by rescuing the inhibitory effect of the guanine-rich 5’ overhangs (**Fig. 7e, f**).

In summary, by modifying the R3C ribozyme substrate with an additional short sequence containing guanine residues, we increased membrane affinity. We had originally hypothesized that enhancing substrate-membrane binding would increase ligation activity by concentrating R3C and its substrate at the membrane surface. The addition of guanines to the substrate, however, introduced an inhibitory effect on ligation activity that was unexpectedly reversed through the introduction of gel-membranes to the reaction. While we did not achieve the intended outcome of enhancing activity, we revealed a lipid-dependent allosteric mechanism for tuning ribozyme activity. These observations raise the possibility that ribozyme activity could have been modulated through lipid-RNA interactions in an RNA World, and provide the proof-of-principle for engineering lipid-sensitive riboswitches with a larger dynamic range for synthetic biology.

## Discussion

Lipids can spontaneously self-assemble to form membranous bilayers, theoretically providing a surface that can concentrate, protect, and regulate RNAs. Here we demonstrate that RNA-gel membrane interactions are dependent on nucleotide content and base pairing, providing a means to engineer RNAs with varying membrane affinities. Increasing the guanine content of the R3C substrate was sufficient to enhance its binding affinity to gel membranes and revealed that ribozyme activity can be regulated in an allosteric lipid-dependent manner. This study yields a framework for engineering RNA-lipid systems that can be regulated based on sequence specificity, and introduces a novel mechanism for riboregulation in cellular and synthetic systems.

Our finding that guanine is a key factor in promoting RNA-gel membrane partitioning is consistent with previous work indicating that guanine residues might play a role in the binding of RNA aptamers to fluid membranes^38^ and that free guanine binds well to fatty acid vesicles^52^. Binding of unstructured oligos containing AG-repeats or deaza-guanine suggests that guanine itself might directly stabilize interactions with the gel membrane, plausibly via hydrogen interactions from the Watson-Crick edge of the nucleotide. It has been proposed that the interaction of adenine and guanine with fatty acid membranes is dependent on the amino group of the nucleotide interacting with carboxyl groups of the fatty acid-based membranes^52^ and that nucleic acid bases can interact with hydrophobic core of the phospholipids^33,35,37^. However, we show in addition that the importance of guanine correlates with its propensity to promote inter- or intra-molecular structures. Guanine can promote structure not only through Watson-Crick base pairs with cytidine, but also with uracil (G-U wobble^57^) and through non-canonical base-paired structures including Hoogsteen base pairs and G-quadruplexes^53^. Indeed, we reveal that G-quadruplexes exhibit enhanced binding to both gel and fluid membranes, providing the impetus to explore whether such interactions are physiologically relevant.

The ability for RNA-lipid interactions to influence ribozyme activity demonstrates a proof-of-principle that spontaneously self-assembling lipid membranes could provide a mechanism for ribo-regulation. In the present study we had hypothesized that gel membranes would enhance R3C ligase activity by increasing concentration of the reactants at the membrane surface. Instead, we discovered that selective guanine-gel membrane interactions had an allosteric effect on the R3C substrate when it was modified with a G-rich tail to enhance membrane binding. This behaviour in some ways resembles the behaviour of riboswitches with regulatory effects that are derived from sensitivity to physicochemical conditions or through binding with an interaction partner^58–62^. However, lipid sensitive riboswitches have not been documented previously. Although the roughly 2-fold change in activity we observe is much smaller than the dynamic range of known riboswitches, it could still be a significant effect in a prebiotic or synthetic system^63^. It is now plausible to explore whether synthetic or naturally occurring RNAs exhibiting sensitivity to lipids could act or be engineered to act as lipid-sensitive riboswitches.

Although gel phase membranes are not widely observed in living cells, RNA-gel membrane interactions could be employed for ribo-regulation in synthetic biological systems. Gel membranes also plausibly accumulated in ancient prebiotic scenarios, potentially serving as an organizational scaffold, as has been proposed for “ribofilms” on mineral surfaces^64^. Thus, RNA-gel membrane interactions provide a plausible means to select RNAs by sequence and structure in primordial and synthetic biological systems. For example, diverse RNA species can be segregated or co-localized based on their differential membrane affinities. Furthermore, selective RNA-membrane localization can be controlled by shifting the temperature above and below the membrane gel-liquid transition temperature, thereby turning on and off RNA-membrane binding. Selective RNA-membrane interactions would also influence the sequence space explored by evolving ribozymes leading to enhanced or novel functions. Looking forward, RNA-gel membrane interactions might facilitate the emergence of novel functions through artificial and natural evolution of ribozymes in the presence of membranes.

In conclusion, our findings reveal that lipids, which are present in every modern cell and were plausibly part of a prebiotic world^45,65^, can interact with RNA and change its activity. These findings have significant applications in fields such as synthetic biology, where merging the selective affinities of aptamers with ribozyme activity (aptazymes) is currently a developing field^67–69^. Furthermore, insights from the present study can already be implemented in bioengineering applications such as in improving mRNA drug delivery mechanisms and introducing lipid sensitive ribo-regulation to synthetic ribozyme networks. More generally, this study gives a simple answer to a fundamental question in the debate on the origin of life – how could primordial RNA molecules be regulated?

## Acknowledgements

We would like to thank Mario Mörl, Gerald Joyce, Ilya Levental, Robert Ernst, Andre Nadler, Dora Tang, Grzegorz Chwastek, Michał Grzybek and Mike Thompson for helpful discussions and feedback. We would also like to thank Anatol Fritsch for home-made microscopy stage-top temperature-control device. This work was supported by the B CUBE of the TU Dresden, a Simons Foundation Fellowship (to J.S.), a German Federal Ministry of Education and Research BMBF grant (to J.S., project 03Z22EN12), and a VW Foundation ‘‘Life’’ grant (to J.S., project 93090).

The authors declare no conflict of interests.

## Materials and methods

### Materials

Dipalmitoylphosphatidylcholine (DPPC), dioleoylphosphatidylcholine (DOPC), dimyristoylphosphatidylcholine (DMPC) and biotinylated phosphatidylethanolamine (18:1 biotinyl cap PE) were purchased from Avanti Polar Lipids (USA) and were used without further purification. Cholesterol was purchased from Sigma Aldrich (USA). DiD was purchased from Invitrogen (USA). HEPES, MgCl_2_ and CaCl_2_ were purchased from CarlRoth (Germany) and were at least >99% pure. All solutions were prepared in MilliQ water, Merck Millipore (USA).

### RNA

The R3C ligase construct^10^ was cloned into the pRZ plasmid^71^ using the InFusion cloning system (Takara Bio, Japan). To ensure correct length of the transcript, pRZ-R3C plasmid was treated with EcoRI-HF (NEB, USA). RNA was expressed using T7 RNA polymerase (homemade, MPI-CBG, Germany) at 37 °C overnight incubation, followed by 10 cycles of 5’ 60 °C -> 5’ 24 °C (HDV cleavage of construct to release pure R3C) and DNAse I treatment (Thermo Scientific, USA). 75-85 nt randomised oligomer was expressed using artificial DNA template (Eurofins Genomics, Germany) – both R3C and randomer were purified by phenol-chloroform-isoamyl alcohol extraction, denaturing PAGE and electroelution.

Synthesis of the deaza-G quadruplex RNA was conducted using T7 transcription on a synthesized DNA template, as described above, however instead of GTP, 7-deaza-GTP (Trilink Biotechnologies, USA) was used. To enhance transcription efficiency, 1 mM GMP was added to the solution^72^.

5’-6-FAM labelled R3C substrates, G quadruplexes as well as other short oligomers used in binding assays were obtained from Integrated DNA Technologies (USA) and used without further purification.

All of the RNA sequences are presented in **Supplementary Table 2**.

### Methods

All of the RNA incubations were prepared in DNA low bind tubes (Sarstedt, Germany) at the constant temperature of 24 °C in buffer composed of 10 mM HEPES pH 7, 5 mM CaCl_2_, and 5 mM MgCl_2_. RNA concentration was determined by measuring absorbance at 260 nm (SPARK 20M, TECAN, Switzerland). Before every incubation RNA was preheated in SafeLock tubes (90 °C, 5’; Eppendorf, Switzerland) to ensure unfolded RNA structures. Denaturing PAGE analysis was performed using 8 – 20% 19:1 acrylamide:bisacrylamide gel composition with 8 M urea, whereas native gels were composed of 6 – 10% 29:1 acrylamide:bisacrylamide mix. Before denaturing PAGE all of the samples were ethanol precipitated.

Structure of R3C ligase from **Fig. 3b** was generated using RNA structure Fold tool from Mathews lab (https://rna.urmc.rochester.edu/RNAstructureWeb/).

### Giant unilamellar vesicle (GUV) preparation and microscopy imaging

Gel-liquid phase separated vesicles were prepared from mixtures of DOPC:DPPC in a 1:1 molar ratio^73^ with 0.5 mol% DiD. 20 nmol of total lipids were evenly distributed on Pt electrodes and dried under a vacuum for 15 minutes. Electroformation (300 Hz, 2.5 V, 65 °C) was conducted in 1 mM HEPES pH 7 and 300 mM sucrose for 3 hours followed by 30’ of lower frequency current to promote GUV formation (2 Hz, 2.5 V, 65 °C).

GUVs were visualized in a home-made microscopy chamber with temperature control in isosmotic solution of glucose and buffer. A Leica DMi8 confocal microscope coupled with a camera (Leica DFC9000 GTC) was used for image acquisition. For both confocal as well as camera acquisition a 63x water-immersed objective was used.

For confocal image acquisition, 488 nm (SybrGold) and 635 nm (DiD) lasers were used at low power (<0.5%) to avoid photobleaching and the signal was collected using hybrid detectors (500 - 550 nm for SybrGold and 650 - 750 nm for DiD). Z-scans were acquired as 4 - 8 averaged images per layer and Z-projected as a sum of the slides using FIJI software^74^.

Temperature ramps were captured using a camera. 1.5 sec of exposure without binning was used to acquire each channel and measurements were repeated every 5 seconds. Temperature changes were registered and calculated to be 3.4 °C/min. Image analysis was conducted in FIJI software.

### Small lipid vesicle preparation

Lipid chloroform stocks were pipetted into a glass vial and briefly evaporated under a steady flow of nitrogen gas. To remove organic solvent residues, a lipid film was dried under vacuum overnight. To obtain multi-layer vesicles (MLVs), lipid films were hydrated in reaction buffer, with a final lipid concentration of 10 mM. Buffer-lipid mixtures were shaken above the melting temperature of the lipids for one hour. A cloudy liposome suspension was freeze-and-thawed 10x and extruded 17x through 100 nm polycarbonate filters (Merck Millipore, USA) to reduce multi-lamellarity and achieve a consistent size distribution of vesicles. Lipid vesicle stocks were kept at 4 °C. Stock lipid concentrations were confirmed using a phosphate assay^75^.

### Dynamic light scattering

The size distribution of lipid vesicles was estimated using a ZetaSizer Nano in 173° backscatter with multiple narrow mode (high resolution) analysis. Final concentration of lipids was 10 - 25 μM and the amount of RNA was fixed at a ratio of 10 lipids / nucleotide, unless stated differently. Results were plotted as the size distribution in the number of detected species (Number PSD).

### UV-crosslinking assay

RNA was incubated for 10 minutes in the reaction buffer with or without lipids followed by 10 minutes incubation under UV-B light (300 mW LEUVA77N50KKU00 LED 305 nm) from a distance of 1.5 – 2 cm. RNA was ethanol precipitated and run on denaturing PAGE. After electrophoresis, gels were poststained with SybrGold dye in order to visualize all of the RNA species. Each crosslinked band was quantified together with whole lane intensity; we define crosslinked bands as band which are higher in mass than the starting RNA and which was not present before UV exposure (**Suppl. 2**). Both values were blanked (subtraction of gel background intensity and “no incubation” sample intensity from the same experimental day). To obtain normalized crosslinking, we’ve divided measured values by the crosslinking efficiency of the lipid-free sample.

### R3C activity assays

5 pmol of R3C ligase was mixed with 0.5 pmol of the substrate to ensure saturation of the system (pseudo 1^st^ order reaction) and to decrease any batch-to-batch differences resulting from R3C purification. Denatured (preheated in 90 °C) RNA was added to the reaction buffer with or without lipid vesicles. Incubation was conducted at various concentrations of lipid vesicles, and samples were ethanol precipitated after 30, 60, 90 and 120 minutes. RNA was analysed on denaturing PAGE using FIJI software. Product and substrate intensities (I_P_ and I_S_, respectively) were quantified and the concentration of product at time t was calculated as:

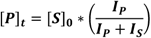

Where:

[P]_t_ – product concentration

[S]_0_ – initial substrate concentration

To determine reaction rates (M/min), all measured product concentration values were fitted using concatenate linear fit with fixed [0;0] intercept. The slope and standard error values of the fit were used for statistical comparisons and data plotting.

### Magnetic beads binding assay

Magnetic beads (DynaBeads streptavidin T1, Invitrogen) were coupled with liposomes doped with 0.5 mol% biotinoyl cap PE. Lipid concentration on beads was estimated using a phosphate assay^75^. Lipid concentration on the non-diluted beads was typically between 500 – 800 μM.

RNA (10 – 25 pmol) was incubated in the buffer and liposome-coated beads for 30’ in 24 °C. As a control (100% samples), non-coupled liposomes were used. After incubation the supernatant was separated from the beads using a magnet. The amount of RNA left in solution was estimated either using absorbance or Qubit miRNA quantification kit (ThermoScientific, USA).

For the absorbance measurements, 20 μL of supernatant was pulled and diluted with 80 μL of MiliQ water and absorbance was measured using 1 cm quartz cuvette. Because of the significant lipid-based light scattering, 100% sample values were calculated from a theoretical approach: knowing RNA concentration we calculated theoretical absorbance in the final RNA dilution.

For the Qubit miRNA quantification assay, 9 μL of supernatant (out of 30 μL reaction mix) was pulled and incubated with 1.5 μL of 0.5 M EDTA (90 °C, 5’). After incubation 150 μl of 1xQubit dye was added to the solution and the amount of RNA was measured using a plate reader (SPARK 20M, TECAN; Ex/Em = 485/530 nm).

The partition coefficient K was calculated according to the following formula:

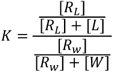

Assuming excess of [L] and [W] over RNA concentration we simplified the equation to:

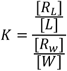

Where:

**[R**_**L**_] – RNA bound to the membranes

**[R**_**W**_] – non-bound RNA

**[L]** – outer lipid concentration

**[W]** – water concentration

[R_W_] was calculated as a ratio or readout from the binding assay sample and the 100% sample. [R_L_] = 1 – [R_W_]. We assumed that RNA interacts only with the outer membrane leaflet, so final outer lipid concentration is equal half of the total lipid amount.

### Ultracentrifugation binding assay

RNA was incubated with various concentrations of DPPC vesicles for 30’. After incubation ultracentrifugation was conducted to pellet vesicles from the solution (125 000 x G, 40’, 24 °C). Measured fluorescence intensity values were plotted as a function of lipid/oligomer or lipid/nucleotide ratios normalized to maximum value in the assay, and fitted using Hill’s fit (OriginLab software):

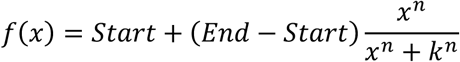

Where Start and End are function plateau values, k is inflection point of curve and n is cooperativity index. The inflection point of the curve (k) and its standard error are parameters which we used for subsequent data analysis.

### Statistical analysis

All of the binding efficiency, partition coefficient, crosslinking efficiency and R3C activity assays were repeated at least 3 times. Error bars in figures represent SEM values. Estimated p values in the figures are result of double sided, unpaired t-student test. We assumed, that a difference is significant if the p value was lower than 0.05.

