## Supplementary figures and tables for "Lipid membranes modulate the activity of RNA through sequence-dependent interactions"

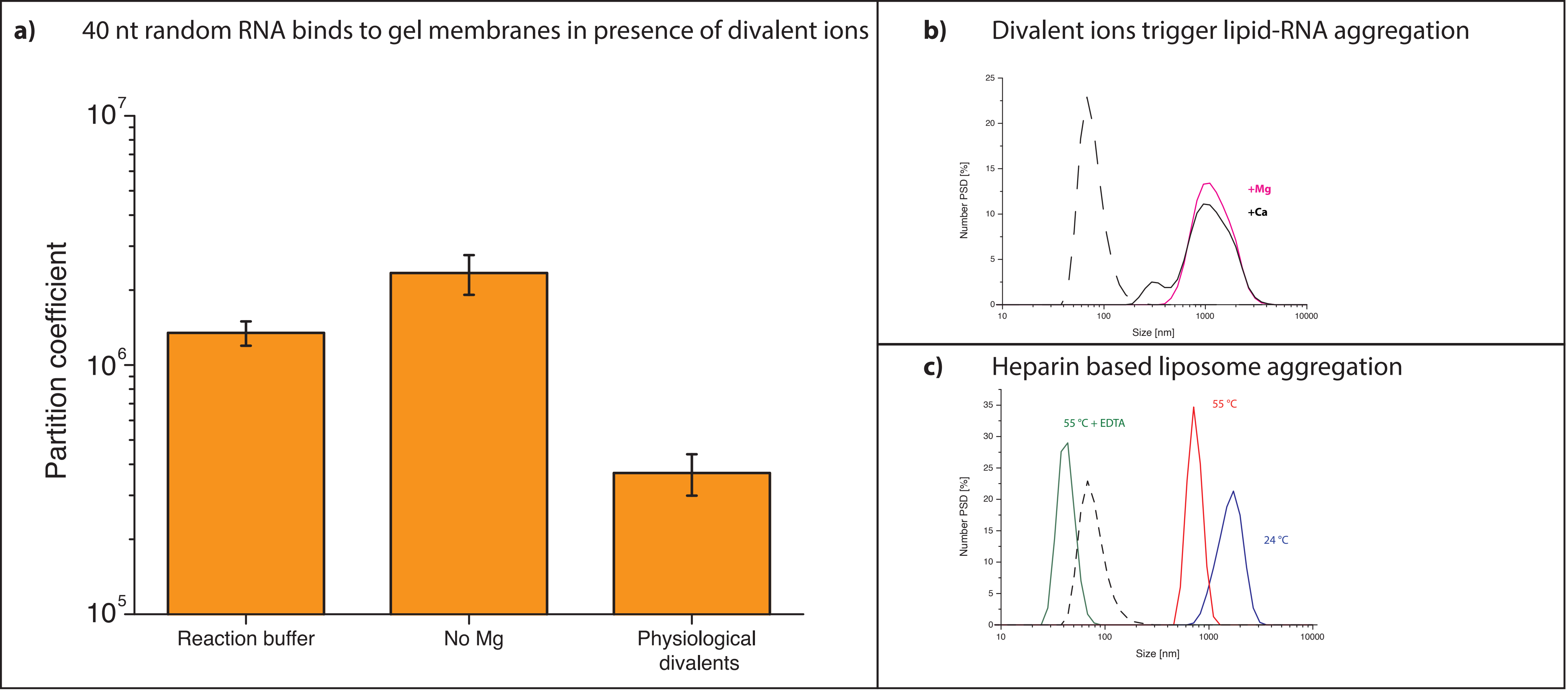


**Supplementary Fig. 1 Both magnesium and calcium can facilitate RNA-lipid interactions. (a)** Depletion of magnesium, which facilitates RNA folding, does not impair RNA randomer-lipid binding. RNA-lipid binding affinity is lower but still well above $10^5$ at physiological concentrations of divalent cations (3 mM $MgCl_2$, 100 nM $CaCl_2$). **(b)** Both magnesium and calcium can induce RNA-based liposome aggregation. Liposomes (25 µM) in the absence of RNA are shown as a dashed line. Final divalent concentration: 5 mM. **(c)** Negatively charged sugar heparin (45 µg/ml) causes significant liposome aggregation suggesting an electrostatic basis of RNA-lipid aggregation. Liposomes (25 µM) in the absence of heparin are presented as dashed line.

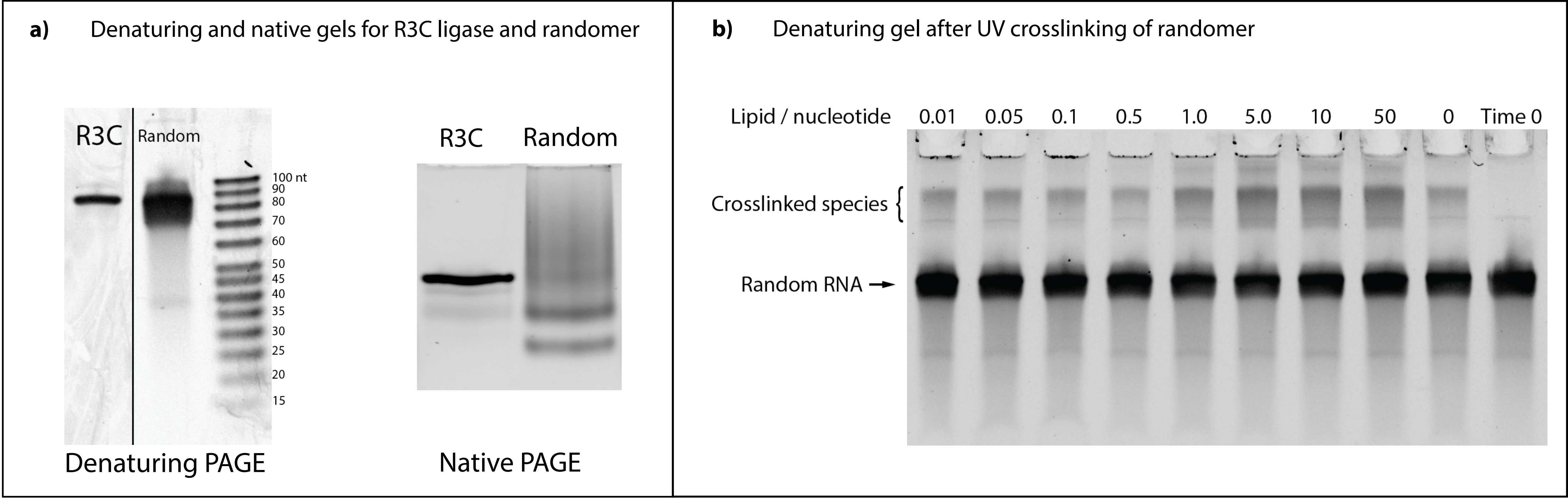


**Supplementary Fig. 2 R3C ligase and randomer size distribution and crosslinking quantification. (a)** The R3C ligase and randomer have a similar size distribution. **(b)** Example of a denaturing gel after UV crosslinking of randomer RNA.

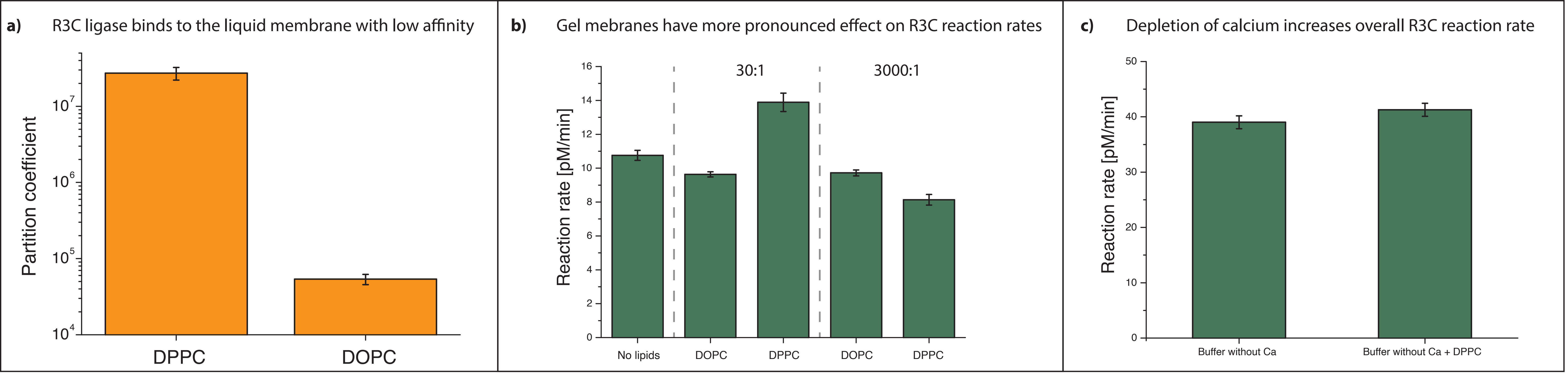


**Supplementary Fig. 3 Gel membranes have larger effect on R3C activity compared with liquid membranes. (a)** R3C ligase binds orders of magnitude better to gel membranes than to liquid membranes. **(b)** R3C ligase reaction rate is not increased in the presence of liquid membranes, in contrast to the reaction in the presence of gel membranes at lipid:R3C = 30 and there is no significant change within broad lipid:R3C ratios (compare lipid:R3C 30:1 and 3000:1, $p$ value > 0.5). **(c)** Depletion of Ca ions causes a 4x increase of R3C reaction rates compared to buffer with Ca ions - see **(b)** reaction in the absence of lipids - however addition of gel membranes does not cause an increase in reaction rate.

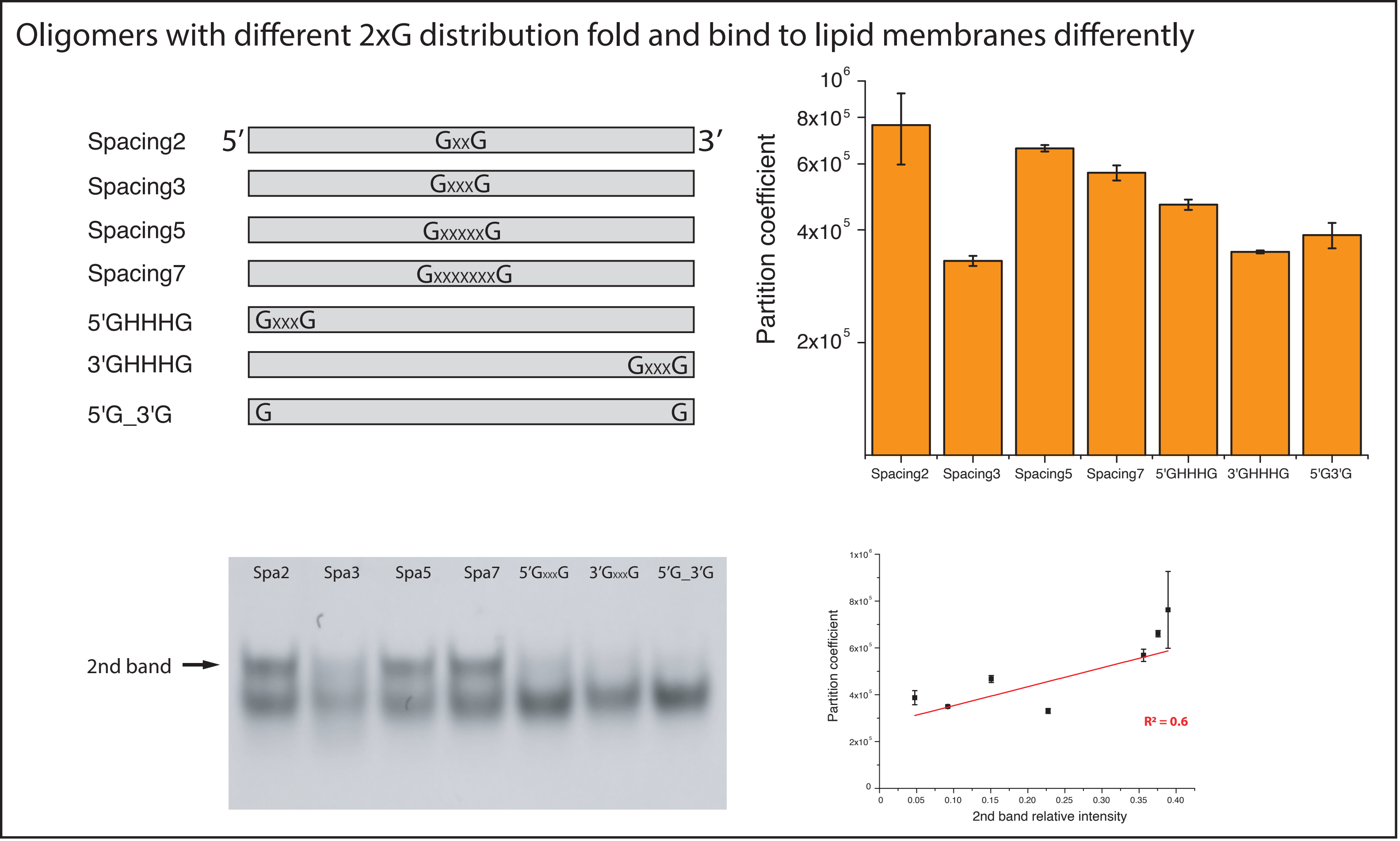


**Supplementary Fig. 4 Presence of the 2nd peak on the native gels for 36 nt oligomers correlates with higher lipid membrane binding.** 2-guanine oligomers show different lipid membrane binding affinities, depending of distribution of the guanines within oligomer. Presence of the RNA structures (2nd peak on native gel) is weakly correlated with lipid binding.

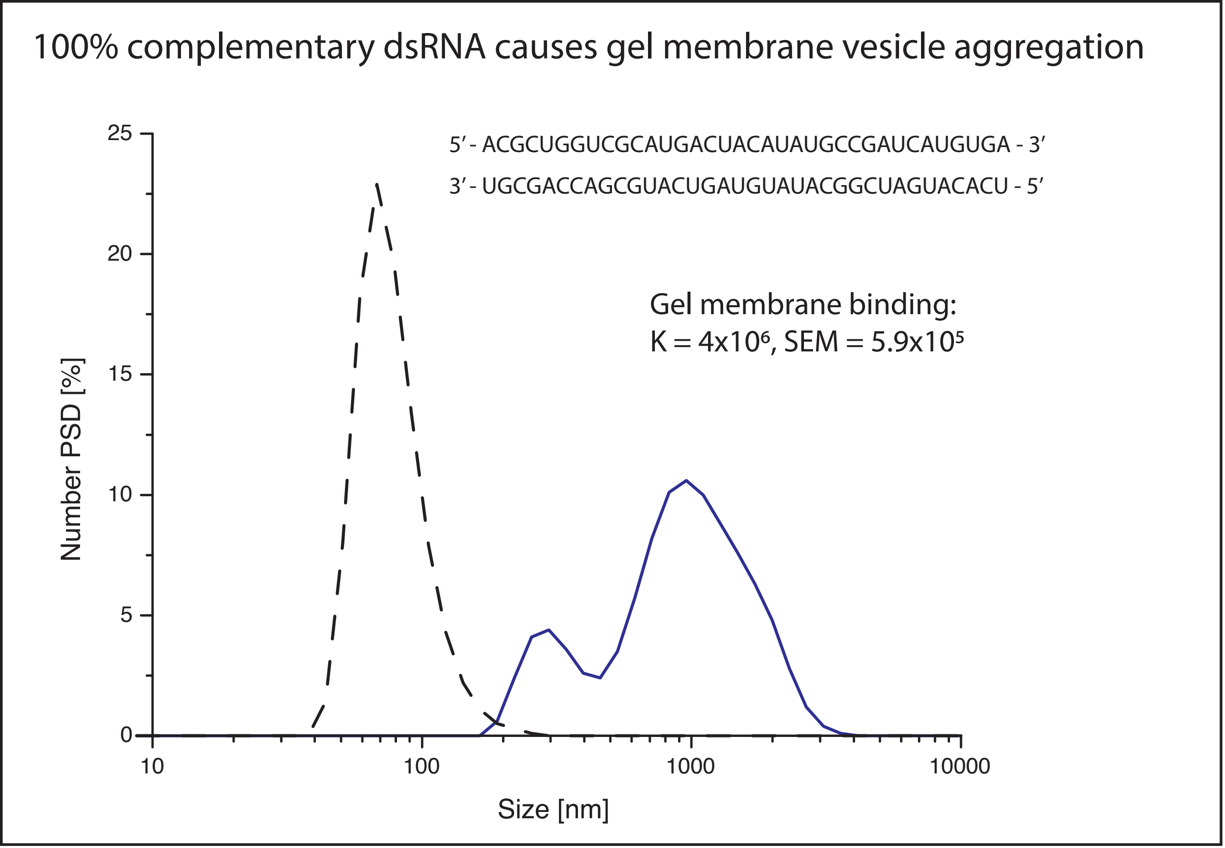


**Supplementary Fig. 5 36 nt double stranded RNA binds to gel membranes and causes vesicle aggregation.** 100% complementary RNA sequences bind with high partition coefficient and promote vesicle aggregation (blue graph). Dashed lines represent DPPC vesicles in the absence of RNA.

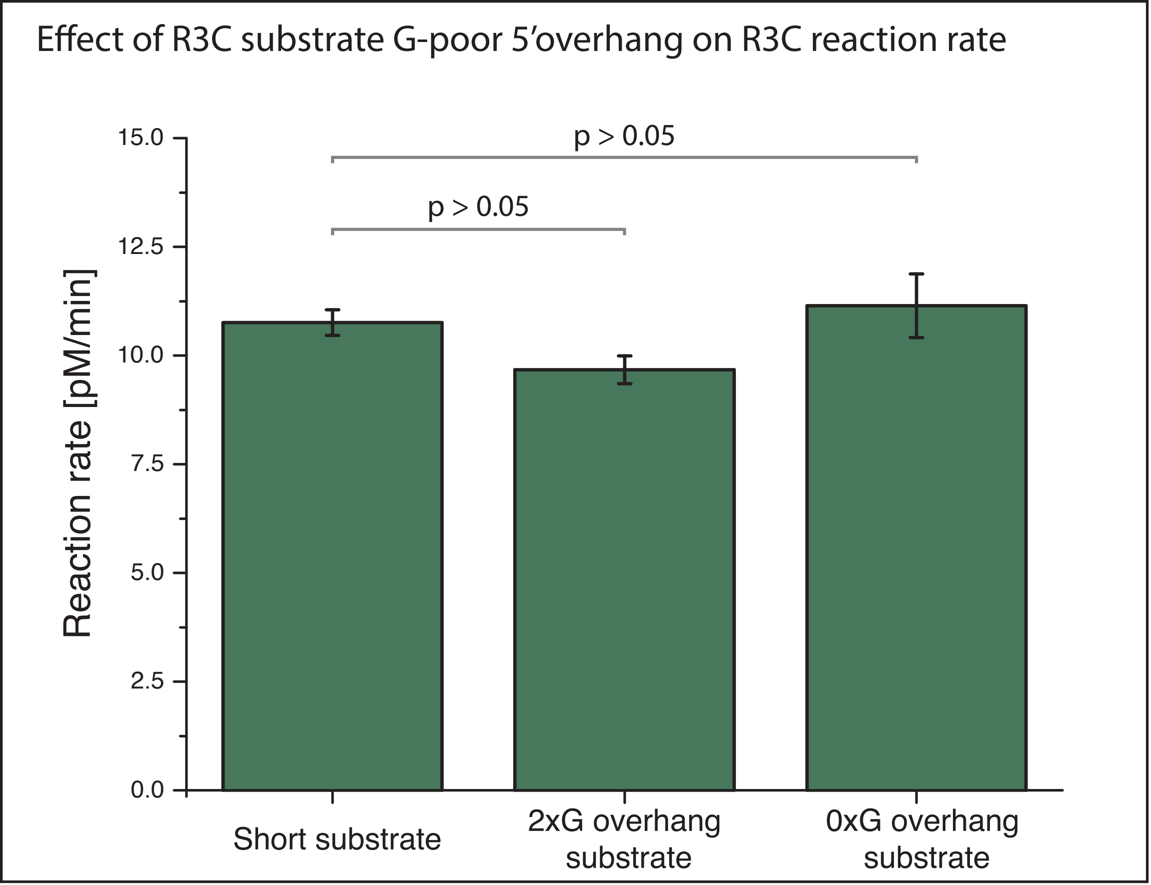


**Supplementary Fig. 6 Modification of R3C substrate with G-deficient 19nt overhang has no effect on activity.** 5’ modification of the R3C substrate with a G-depleted randomized 19 nt overhang had no significant effect on ligase activity. Furthermore, modification with a 2xG randomized 19 nt overhang produced a relatively small effect.

| Ribozyme | Type of activity | Plausible feature of binding to the lipid membrane | Ref. |
|---|---|---|---|
| HDV | Cis, cleavage | • Stabilisation/destabilisation of pseudoknot structure can lead to increase/decrease of the ribozymatically active structures | Ferré-D'Amaré et al. 1998 |
| Hammerhead | Trans, cleavage | • Upconcentration on the membrane surface of substrate and ribozyme<br>• Retaining longer RNA species on the membrane surfaces - switching reaction equilibrium<br>• Stabilisation/destabilisation of substrate-ribozyme structure | Drobot et al. 2018 |
| Ligase (R3C)<br>Replicase | Trans, ligation | • Stabilisation/destabilisation of substrate-ribozyme structure<br>• Upconcentration on the membrane surface of substrate and ribozyme | Rogers et al. 2001<br>Lincoln et al. 2009 |
| Polymerase | Trans, condensation | • Stabilisation/destabilisation of ribozyme structure<br>• Upconcentration of the reaction substrates on the membrane surface | Müller et al. 2008 |

**Suppl. Table 1 – Possible effects of RNA-lipid interactions on different types of ribozymes.** Various ribozyme species could either gain or lose functionality upon membrane binding. For example, the Cis acting HDV ribozyme might be biased by changes to the catalytically crucial pseudoknot structure. Trans acting ribozymes in general can benefit from colocalization and up-concentration of ribozyme and substrates on the membrane, which can enhance reaction rate. Hammerhead as a multiple turnover ribozyme could potentially benefit from separation of the shorter cleavage products from the longer substrate due to selective retention of longer RNAs at the membrane surface, which can push the reaction equilibrium toward more product. R3C ligase and replicase based on R3C as single turnover trans acting ribozyme can benefit only from up-concentration of the reactants and possibly through structural effects resulting from RNA-lipid binding such as stabilization of reaction intermediates. Lastly, polymerase ribozymes would benefit mostly from up-concentration at the membrane surface.

Ferré-D'Amaré, A. R., Zhou, K. & Doudna, J. A. Crystal structure of a hepatitis delta virus ribozyme. Nature 395, 567–574 (1998).

Drobot, B., Iglesias-Artola, J. M., Vay, K. L., Mayr, V., Kar, M., Kreysing, M., Mutschler, H. & Tang, T.-Y. D. Compartmentalised RNA catalysis in membrane-free coacervate protocells. Nat Commun 9, 3643 (2018).

Rogers, J. & Joyce, G. F. The effect of cytidine on the structure and function of an RNA ligase ribozyme. Rna 7, 395–404 (2001).

Lincoln, T. A. & Joyce, G. F. Self-Sustained Replication of an RNA Enzyme. Science 323, 1229–1232 (2009).

Müller, U. F. & Bartel, D. P. Improved polymerase ribozyme efficiency on hydrophobic assemblies. Rna 14, 552–562 (2008).

Supplementary table 2 – used RNA sequences

| Sequence name | Sequence (5' – 3') |
|---|---|
| **R3C** | GAGACCGUAAUGAGUAGUACUUAUUAUGCUCGAUUGUUCGUAA GAACAGUUUGAAUGGGUUGAAUAUAGUGAGGAUCCGCUAGC |
| **~80 nt randomer** | 75-85xN |
| **Short R3C substrate** | 6_FAM_CGACUCACUAUA |
| **4xAG substrate** | 6_FAM_A**G**A**G**A**G**A**G**CGACUCACUAUA |
| **9xAG substrate** | 6_FAM_A**G**A**G**A**G**A**G**A**G**A**G**A**G**A**G**A**G**ACGACUCACUAUA |
| **2G substrate** | 6_FAM_HHHHHHHH**G**HHH**G**HHHHHHCGACUCACUAUA |
| **0G substrate** | 6_FAM_HHHHHHHHHHHHHHHHHHHCGACUCACUAUA |
| **9xUC for 9xAG screening** | UCUCUCUCUCUCUCUCUCU |
| **0G** | HHHHHHHHHHHHHHHHHHHHHHHHHHHHHHHHHHHH |
| **2G** | HHHHHHHHHHHHHHHH**G**HHH**G**HHHHHHHHHHHHHHHH |
| **4G** | HHHHHHHHHHHH**G**HHH**G**HHH**G**HHH**G**HHHHHHHHHHH |
| **5G** | HHHHHHHHHH**G**HHH**G**HHH**G**HHH**G**HHH**G**HHHHHHHHHH |
| **7G** | HHHHHH**G**HHH**G**HHH**G**HHH**G**HHH**G**HHH**G**HHH**G**HHHHHH |
| **9G** | HH**G**HHH**G**HHH**G**HHH**G**HHH**G**HHH**G**HHH**G**HHH**G**HHH**G**HH |
| **2G_spacing 2** | HHHHHHHHHHHHHHHH**G**HH**G**HHHHHHHHHHHHHHHH |
| **2G_spacing 5** | HHHHHHHHHHHHHH**G**HHHHH**G**HHHHHHHHHHHHHHH |
| **2G_spacing 7** | HHHHHHHHHHHHH**G**HHHHHHH**G**HHHHHHHHHHHHHH |
| **2G_5'end** | **G**HHH**G**HHHHHHHHHHHHHHHHHHHHHHHHHHHHHHH |
| **2G_3'end** | HHHHHHHHHHHHHHHHHHHHHHHHHHHHHHH**G**HHH**G** |
| **2G_5'G_3'G** | **G**HHHHHHHHHHHHHHHHHHHHHHHHHHHHHHHHHH**G** |
| **40xN** | NNNNNNNNNNNNNNNNNNNNNNNNNNNNNNNNNNNNNNNN |
| **40xN - G** | HHHHHHHHHHHHHHHHHHHHHHHHHHHHHHHHHHHHHHHH |
| **40xN - A** | BBBBBBBBBBBBBBBBBBBBBBBBBBBBBBBBBBBBBBBB |
| **40xN - C** | DDDDDDDDDDDDDDDDDDDDDDDDDDDDDDDDDDDDDDDD |
| **40xN - U** | VVVVVVVVVVVVVVVVVVVVVVVVVVVVVVVVVVVVVVVV |
| **19xG** | **GGGGGGGGGGGGGGGGGGG** |
| **19xA** | AAAAAAAAAAAAAAAAAAA |
| **19xC** | CCCCCCCCCCCCCCCCCCC |
| **19xU** | UUUUUUUUUUUUUUUUUUU |
| **19xN** | NNNNNNNNNNNNNNNNNNN |
| **AG repeats** | 6_FAM_AGAGAGAGAGAGAGAGAGA |
| **ssRNA (ACUG)** | ACUGACUGACUGACUGACUGACUGACUGACUG |
| **ssRNA (CAGU)** | CAGUCAGUCAGUCAGUCAGUCAGUCAGUCAGU |
| **dsRNA, not overlapping** | ACGCUGGUCGCAUGACUACAUAUGCCGAUCAUGUGA premixed with UCACAUGAUCGGCAUAUGUAGUCAUGCGACCAGCGU |
| **Hairpin RNA** | GCCGAUCACAUGACCAUAUAUGGUCAUGUGAUCGGC |

| | |
|---|---|
| **G quadruplex** | 6_FAM_AGGGAAGGGAAGGGAAGGG |
| **Deaza-G quadruplex** | GggAAgggAAgggAAgggA |

**B** – C, G or U
**D** – A, G or U
**H** – A, C or U
**V** – A, C or G
**g** – 7-deaza-guanine
